# CSB-dependent CDK9 degradation and RNA Polymerase II phosphorylation during Transcription Coupled Repair

**DOI:** 10.1101/316935

**Authors:** Lise-Marie Donnio, Anna Lagarou, Gabrielle Sueur, Pierre-Olivier Mari, Giuseppina Giglia-Mari

**Author notes:** to whom correspondence and request for materials should be addressed: G G-M.

## Abstract

DNA lesions block cellular processes such as transcription, inducing apoptosis, tissue failures and premature ageing. To counteract the deleterious effects of DNA damage, cells are equipped with various DNA repair pathways. Transcription Coupled Repair specifically removes helix-distorting DNA adducts in a coordinated multi-step process. This process has been extensively studied, however once the repair reaction is accomplished, little is known about how transcription restarts. In this study, we show that, after UV irradiation, the CDK9/CyclinT1 kinase unit is specifically released from the HEXIM1 complex and that this released fraction is degraded in the absence of CSB. We determine that UV-irradiation induces a specific Ser2 phosphorylation of the RNA polymerase II and that this phosphorylation is CSB dependent. Surprisingly CDK9 is not responsible for this phosphorylation but instead plays a non-enzymatic role in transcription restart after DNA repair.

## Introduction

Cells are the units of organic life and store in their nuclei, under the form of the DNA molecule, what constitutes the instruction manual for proper cellular functioning. Despite the protection offered by the cellular environment, the integrity of DNA is continuously challenged by a variety of endogenous and exogenous agents (*e.g.* ultraviolet light, cigarette smoke, environmental pollution, oxidative damage, etc…) that cause DNA lesions, interfering with proper cellular functions, *in fine* causing the aging or premature aging of the tissue and later on of the whole organism.

To prevent the deleterious consequences of persisting DNA lesions, all organisms are equipped with a network of efficient DNA damage responses and DNA repair systems. One of these systems is the <u>N</u>ucleotide <u>E</u>xcision <u>R</u>epair (NER). NER removes helix-distorting DNA adducts such as UV-induced lesions (Cyclo-Pyrimidine Dimers and 6-4 Photoproducts, CPD and 6-4PP) in a coordinated multi-step process (1).

The NER system has been linked to rare human diseases classically grouped into three distinct NER-related syndromes. These include the highly cancer prone disorder xeroderma pigmentosum (XP) and the two progeroid diseases: Cockayne syndrome (CS) and trichothiodystrophy (TTD) (2). Importantly, CS and TTD patients are not cancer-prone but present severe neurological and developmental features.

NER exists in two distinct sub-pathways depending where DNA lesions are located within the genome. <u>G</u>lobal <u>G</u>enome <u>R</u>epair (GG-NER or GGR) will repair DNA lesion located on non-transcribed DNA. While, the second sub-pathway is directly coupled to transcription elongation and repairs DNA lesions located on the transcribed strand of active genes and it is designated as <u>T</u>ranscription-<u>C</u>oupled <u>R</u>epair (TC-NER or TCR).

RNAP2 frequently deals with obstacles that need to be removed through the TCR process for resumption of transcription (3). Constant blockage of transcription has severe consequences for the cell, since it might be a signal for apoptosis. Deficient TCR is illustrated in CS patients, a rare inherited syndrome characterized by multi-system clinical malfunctions, growth and neurological abnormalities and features of premature ageing due to increased apoptosis. At the cellular level, a hallmark of CS is the inability to resume RNA synthesis after exposure to UV-light (4-6). This not only identifies TCR as a crucial defense mechanism against DNA damage for cells and organisms to escape from lethal effects of transcription inhibition, but also highlights the great importance of transcriptional resumption *after* repair of the damaged transcribed strand.

During a TCR event two phases can be distinguished: (i) the actual repair of the damaged strand *via* the TCR sub-pathway and (ii) the <u>r</u>esumption of <u>t</u>ranscription after <u>r</u>epair (RTR).

Although the TCR repair process has been extensively described, the molecular mechanisms implicated in RTR and the specific proteins involved are still elusive. The regulation of resumption of transcription after repair is highly important given that improper restart leads to cellular malfunction and apoptosis, and concomitantly contributes to ageing.

Interestingly, there has been some recent progress, concerning the complex, yet poorly defined, mechanism, which allows transcription resumption after DNA repair. These studies open the way for a deeper understanding on the RTR mechanism at different levels (7) (8) (9) (10). One of these studies describe the identification of an RNAP2 elongation factor (ELL: eleven-nineteen lysine-rich leukemia) as a new partner of the basal transcription repair factor TFIIH (8). The best-characterized function of ELL is to increase the catalytic rate of RNA Polymerase II transcription by suppressing transient pausing by Polymerase at multiple sites along the DNA during elongation (11). The combination of the UV-sensitivity, the absence of RNA recovery synthesis (RRS) and the unchanged DNA synthesis (UDS), illustrated in ELL-depleted cells upon UV-irradiation, suggest that ELL is an indirect TCR-repair factor, which plays a more specific role during RTR. To date, these results favor a possible model where ELL is recruited to the arrested RNAP2 by its interaction with TFIIH and functions as a platform for other elongation factors in order to facilitate RTR (8).

Several groups have reported *in vitro* that ELL and the positive transcription elongation factor b (P-TEFb) exist in complexes with multiple MLL translocation partners, called Super Elongation Complexes (SECs) (12). P-TEFb consists of a heterodimeric kinase, composed of CDK9 and its Cyclins (K/T1/T2), which play a central role in the release of RNAP2 from pausing. In mammalian cells, the CDK9 subunit of pTEF-b phosphorylates RNAP2 at its Ser-2 carboxy-terminal domain (CTD) repeat to license assembly of multiple factors critical for mRNA biogenesis, chromatin modification during transcription (13) and for marking transcription elongation (14). Interestingly, phosphorylation of the CTD does not directly affect elongation rate but instead mediates interactions between the polymerase and other factors (15). Therefore, the Ser5P to Ser2P transition could either promote the association and activity of positive elongation factors, or inhibit pathways that causes RNAP2 to pause or terminate early in elongation. Hence it is possible that ELL functions via its interaction with the positive elongation factor (pTEF-b) and its subsequent recruitment, in order to facilitate transcriptional resumption after repair of the transcription blocking DNA lesion.

In this article we investigated the role of CDK9 during transcription-coupled repair (TCR) and most precisely during resumption of transcription after DNA repair (RTR). Our results clearly show that CDK9 plays a role during RTR but that, differently from its function during release of paused RNAP2 molecules, this role in not dependent on its kinase activity and that, in the absence of CDK9, RTR is delayed and RNAP2 molecules remain bound to the chromatin after UV irradiation. Interestingly, we could highlight a specific DNA-damage dependent dissociation of CDK9 and CycT from the 7SK snRNP complex and a degradation of CDK9 and CycT after UV-irradiation in TCR deficient cells. Finally, we could reveal that during RTR the RNAP2 is unambiguously phosphorylated at the Ser2 position (independently from Cdk9 kinase activity) and that this phosphorylation is absent in TCR-deficient cells.

## Results

### CDK9 function during Nucleotide Excision Repair

Because of the strong relation between ELL and CDK9 (12), we decided to explore the role of CDK9 in DNA Repair and particularly within the Nucleotide Excision Repair system. This repair pathway can be subdivided into GGR (Global Genome Repair) and TCR (Transcription Couple Repair).

Repressing CDK9 expression by siRNA in normal human MRC5 fibroblasts led to no UV-specific cytotoxicity, comparable to mock-knocked down cells, as measured by clonogenic survival (Fig 1A). As a positive control, the knockdown of the well-known NER endonuclease XPF was used, which, as expected, produced severe UV cytotoxicity. This result clearly indicates that CDK9 absence does not confer UV sensitivity to cells.

**Figure 1.**
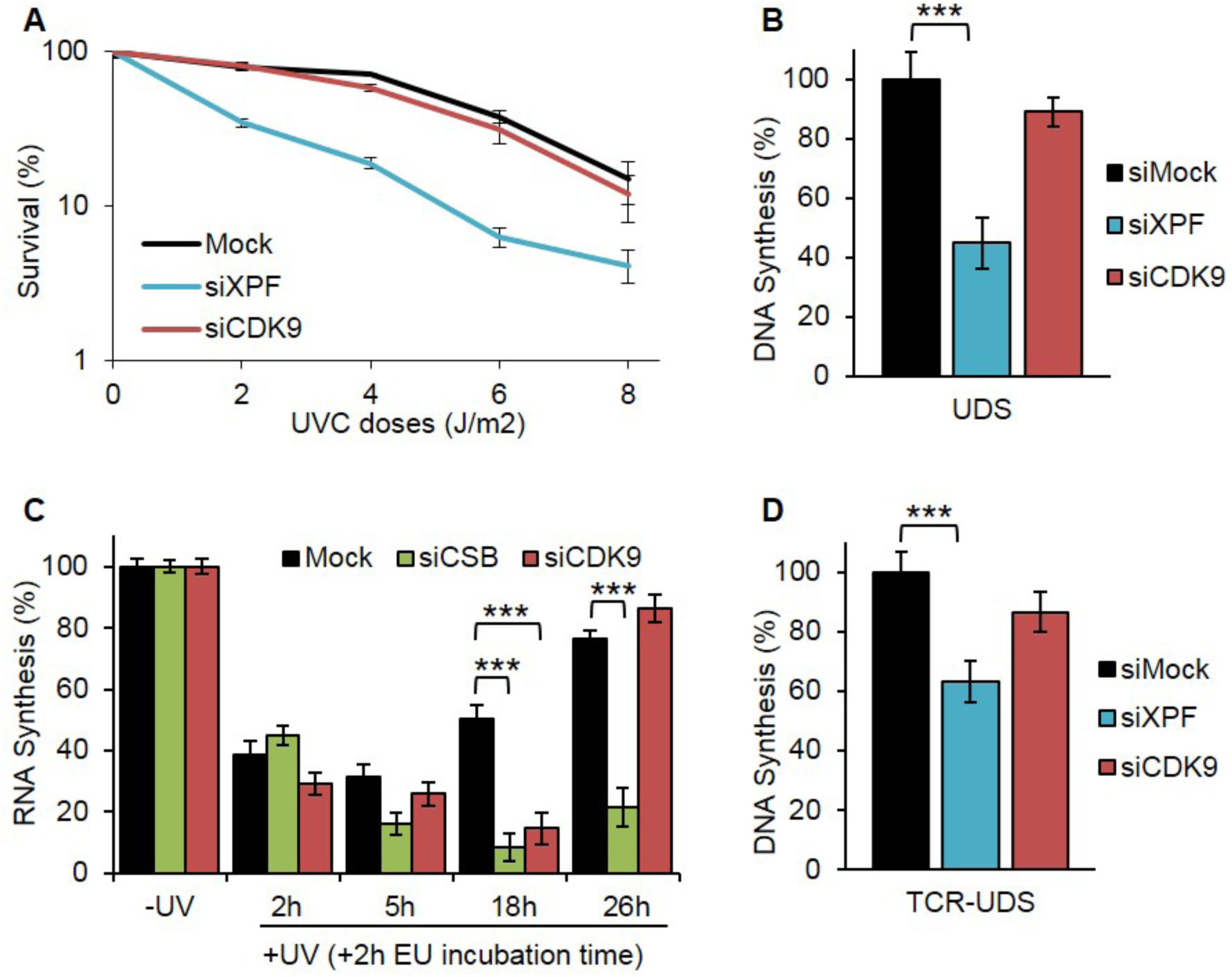
CDK9 function in NER. **A** Sensitivity to **UV-C** of immortalized human fibroblast MRC5 cells treated with siRNA against the indicated factors as determined by colony-forming ability. **B** UDS determined by EdU incorporation after local UV-C exposure in MRC5 cells after siRNA mediated knockdown of the indicated factors. At least 20 nuclei were analyzed. **C** RRS after UV-C exposure in MRC5 cells after siRNA mediated knockdown of the indicated factors. At least 60 nuclei were analysed. **D** TCR-UDS determined by EdU incorporation after local UV-C exposure in a GGR-deficient cell line (XPC-/-) after siRNA mediated knockdown of the indicated factors. At least 25 local damages were analysed. For all figures, mock siRNA (black), XPF siRNA (blue), CDK9 siRNA (red), CSB siRNA (green); error bars represent the SEM. p-value: ^***^<0,001

While cell survival after UV treatment is a general measure of the cellular DNA repair activity, the golden standard to quantify GGR activity is the Unscheduled DNA Synthesis after UV treatment. This assay quantifies DNA replication after repair, i.e. the refilling of single-strand DNA gaps generated by NER processing of UV-induced DNA lesions within locally UV-exposed cell nuclei. As expected for NER-deficient cells unable to process UV lesions, XPF siRNA treated cells showed a strong reduction in UDS levels (Fig 1B). CDK9-depleted cells showed no significant decrease in UDS levels, as in mock-knock down cells, suggesting that CDK9 is not an essential factor during GGR. Additionally, no endogenous CDK9 local damage accumulation was observed when MRC5 cells where locally UV-irradiated (Fig S1).

These results show that CDK9 is not involved in the GGR sub-pathway but do not exclude that CDK9, as hypothesize by its role within transcription, could be involved in TCR and RTR. In fact, proteins involved in TCR are not easily visualized on the damage areas. This is mainly due to the low level of TCR (10%) versus GGR (90%) in the cells (8) and intrinsically because the TCR reactions are primarily related to the transcribed genes, their number is hence limited to the number of active transcriptions.

The golden standard assay to measure TCR activity in the cell is the RRS (<u>R</u>NA-<u>R</u>ecovery <u>S</u>ynthesis). In this assay, transcriptional activity is visualized by detecting and quantifying the newly synthesized RNA via the incorporation of a fluorophore-coupled nucleoside analog. The experiment is conducted at different time points after UV induction (2h, 5h, 18h and 26 h), allowing quantifying the decline in transcriptional activity after UV damage (5 hours after UV irradiation) and the restart of activity after repair of the DNA damage (18 hours after UV irradiation). We used this test to quantify TCR activity in globally UV-irradiated siRNA-treated cells. Because CDK9 silencing might affect global transcription (16), we quantified the residual transcriptional activity during our experiments. We could measure that 30% CDK9 protein is still present in the cells after silencing (Fig S2) and that this amount results in a 70% residual basal transcription activity (Fig S3). In order to quantify only the effect of CDK9 silencing on TCR, we took into account the reduction in transcriptional activity observed when CDK9 is silenced and normalized the mRNA production at 2, 5, 18 and 26 hours to the non-irradiated condition. As a positive control during RRS, CSB (an established repair factor, working specifically in TCR) knocked-down cells have been used. As expected, in the absence of CSB, no restart of transcription after UV damage is observed. Knockdown of CDK9 resulted in a significantly reduced RNA synthesis at 18 h after UV with a full recovery of RNA synthesis at later time points (Fig 1C), showing that the reduction of CDK9 retards the recovery of transcription after UV damage, affecting the TCR pathway (either retarding the repair reaction or delaying the restart of transcription after completion of DNA repair).

In order to discriminate whether CDK9 plays a role in the repair process or in the restart of transcription after repair, we performed an assay designed previously in our group which specifically measures repair replication during TCR: the TCR-UDS assay (8). We treated with siCDK9 and siXPF, XPC-mutant cells (GGR-deficient) in order to be able to monitor only TCR-specific replication levels within locally UV-exposed nuclear regions. In order to precisely localize the UV-induced DNA-damaged areas, a co-immunofluorescence labeling of γ-H2AX was performed and repair replication was quantified. In siXPF treated XPC-negative cells both GGR and TCR pathways are compromised and, as expected, low TCR-UDS levels were observed (Fig 1D). In contrast, the knockdown of CDK9 resulted in normal TCR-UDS levels comparable to the levels seen in mock-knocked down cells. This result combined with the RRS results (Fig 1C) shows that CDK9 is not involved in the repair reaction *per se*, but that <u>R</u>esumption of <u>T</u>ranscription after DNA <u>R</u>epair (RTR) is delayed when CDK9 is knocked down.

### UV-irradiation induces a dissociation of the CDK9-HEXIM1 complex

In order to study in details the role of CDK9 during RTR, we generated stably expressing GFP-tagged CDK9 (CDK9-GFP) SV40-immortalized human fibroblasts (MRC5SV40, here after MRC5). A simplified scheme of the recombinant fused-protein is depicted in Figure S4A. High-resolution confocal imaging of these cell lines revealed that CDK9-GFP is mainly present in the nucleoplasm, and absent in the nucleoli and/or the cytoplasm (Fig S4B). By performing immunofluorescence experiments on fixed-cells exogenously expressing CDK9-GFP, we confirmed that the cellular localization of the recombinant protein is very similar to that of the endogenous proteins (Fig S4B). Immunoblot analysis on whole cell extracts of CDK9-GFP expressing cells has been used to quantify the ratio of the recombinant protein expression in comparison with the endogenous untagged protein. As shown in Figure S4C, CDK9-GFP expression is equivalent to the endogenous CDK9.

We could previously show that ELL is localized on locally damaged nuclear regions (8). Because of the physical interactions between ELL and CDK9 (12) we investigated whether CDK9 is also recruited on these damaged regions. We compared CDK9 immediate recruitment on DNA lesions to the accumulation of the TCR-specific protein CSA (Fig 2A). Confocal time-lapse images of living CDK9-GFP and CSA-GFP were taken after induction of local damage and the accumulation of the two proteins on these locally damaged areas was quantified. Interestingly, in contrast to CSA and ELL (8), there is no accumulation of CDK9 to the DNA damage. (Fig2A and 2B). This is not due to higher expression of the endogenous CDK9 in comparison with the GFP-tagged version, since their expression is equivalent as shown above (Fig S4C).

**Figure 2.**
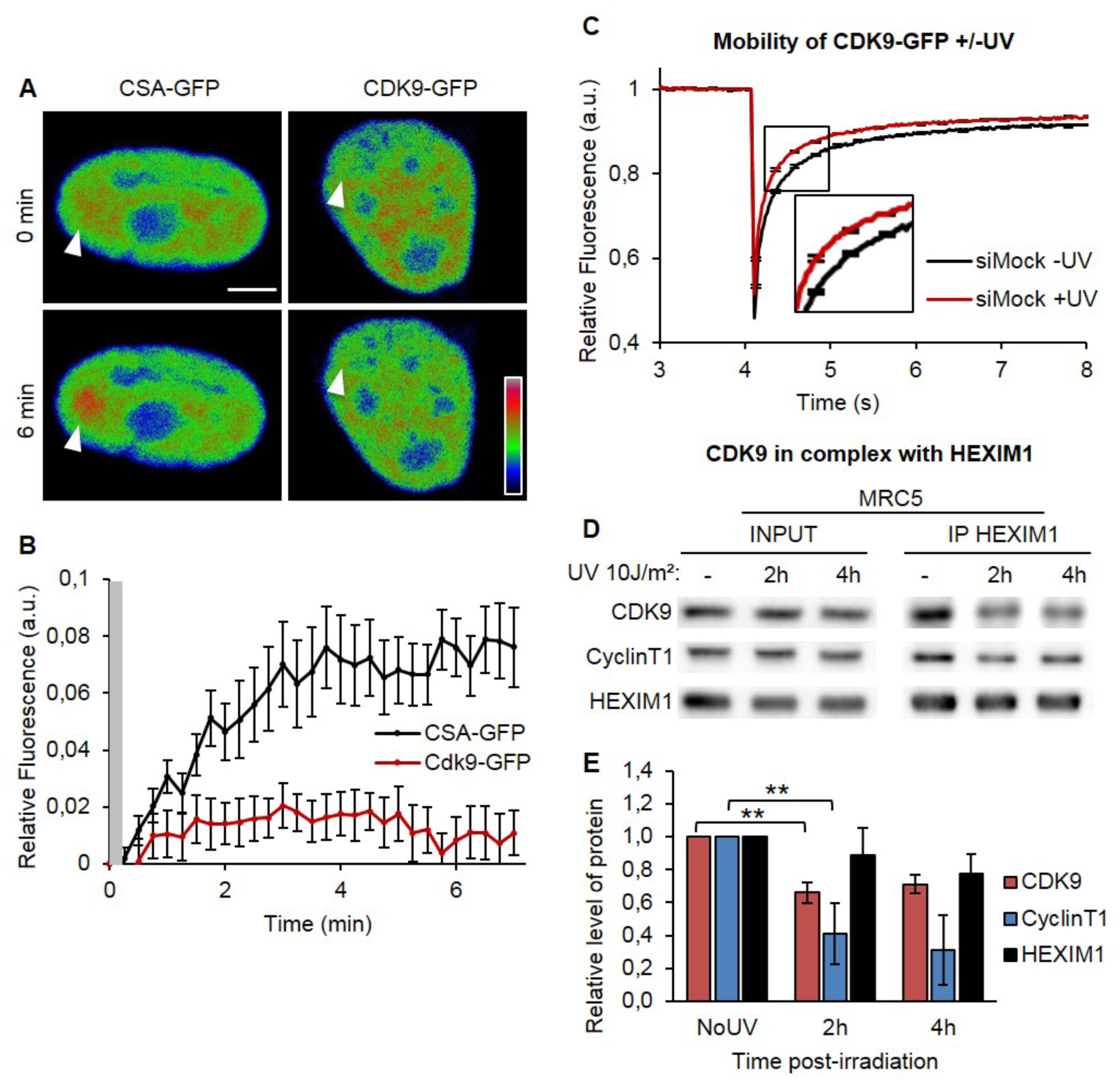
CDK9 dynamics during DNA damage response. **A** Confocal time-lapse images of a living mammalian fibroblast expressing CSA-GFP or CDK9-GFP, seen accumulating at a microirradiated area (arrows). Images with rainbow RGB Lookup table. Scale bar = 5μm. **B** Accumulation curves of CDK9-GFP (red line) and CSA-GFP (black line) proteins at laser-induced DNA damage. Hatched region indicates the time window during which microirradiation-induced photobleaching masks the accumulation signal. Error bars represent the SEM obtained from at least 14 cells. **C** Normalized FRAP analysis of CDK9-GFP expressing cells after siRNA mediated knockdown of the indicated factors. Cells were measured untreated (-UV) or 4h after UC-C exposure (+UV). Error bars represent the SEM obtained from at least 27 cells from 2 independent experiments. p-value between the two curves inferior to 0,001. **D** Immunoprecipitation of HEXIM1 in MRC5 cells. Bound proteins were revealed by Western blot using antibodies against CyclinT1, CDK9 and HEXIM1. INPUT corresponds to 20% of the lysate used for IP reactions. **E** Quantification of at least three different experiments of the IP/INPUT ratio compared with NoUV condition. p-value: ^**^<0,05

However, it has to be noticed that these recruitment measurements were obtained seconds after the induction of DNA damage. To quantify a possible recruitment of CDK9 on damaged DNA at later time points after UV irradiation, we applied the Strip-FRAP (<u>F</u>luorescence <u>R</u>ecovery <u>A</u>fter <u>P</u>hotobleaching) method (17), in which fluorescent molecules are photo-bleached in a small strip by a high intensity laser pulse and then the subsequent recovery of fluorescence is monitored in time within the bleached area. Without any damage, this measure of fluorescence recovery corresponds to the protein mobility within the living cell. However, when UV-irradiation is applied, a protein that physically interacts with the damage will be slowed down (due to the interactions with the substrate) and the recovery of fluorescence will be reduced. Unexpectedly, as shown in Figure 2C, instead of having a reduced mobility (as repair proteins have), CDK9 becomes more mobile after UV irradiation. This might suggest the existence of fraction of CDK9 that is part of a larger complex with lower mobility, from which CDK9 dissociates upon UV-induced DNA damage in order to execute its specific role during the RTR process. In absence of damage, CDK9 interacts physically with HEXIM1 (18), a subunit of the 7SK snRNP complex, which inhibits the kinase activity of CDK9 (18) within the P-TEFb complex. Because UV-dependent transcription inhibition induces a dissociation of P-TEFb from the 7SK snRNP complex (19), we wanted to confirm that in our experiments and at our time points, CDK9 increased mobility, measured by FRAP, could be a result of the dissociation of CDK9 from HEXIM1. In order to verify this hypothesis, we immuno-precipitated HEXIM1 and quantified the amount of CDK9 and CyclinT1 found together with HEXIM1. As expected, in absence of DNA damage, HEXIM1 and CDK9/CyclinT1 could be co-immuno-precipitated (Fig 2D and 2E). However, 2 to 4 hours after UV-irradiation a lower amount of CDK9/CyclinT1 could be immuno-precipitated together with HEXIM1 showing, as previously reported (19) that UV-irradiation causes a dissociation of CDK9/CyclinT1 from HEXIM1 (Fig 2D and 2E), demonstrating that our FRAP method can be used to measure CDK9/HEXIM1 UV-dependent dissociation.

### UV-dependent degradation of CDK9/Cyclin 1 in the absence of CSB

To verify that CDK9/HEXIM1 UV-dependent dissociation is also dependent on the TCR reaction, we measure CDK9 mobility upon UV damage in the absence of CSB, hence a TCR deficient background. Interestingly, in the absence of CSB, the release of CDK9 from HEXIM1 upon UV-induction is lost (Fig 3A and Fig2C). This result can be explained by invocating that the remobilization from HEXIM1 of CDK9 after UV damage takes place via a CSB and probably TCR-dependent mechanism or that the released fraction is rapidly degraded in the absence of CSB. This last hypothesis was confirmed by the fact that unexpectedly, we could notice that after UV-irradiation, in both CSB deficient cells and CSB knocked down cells, the amount of CDK9/CyclinT1 was consistently reduced compared to the amount of HEXIM1 (Fig 3B and 3C).

**Figure 3.**
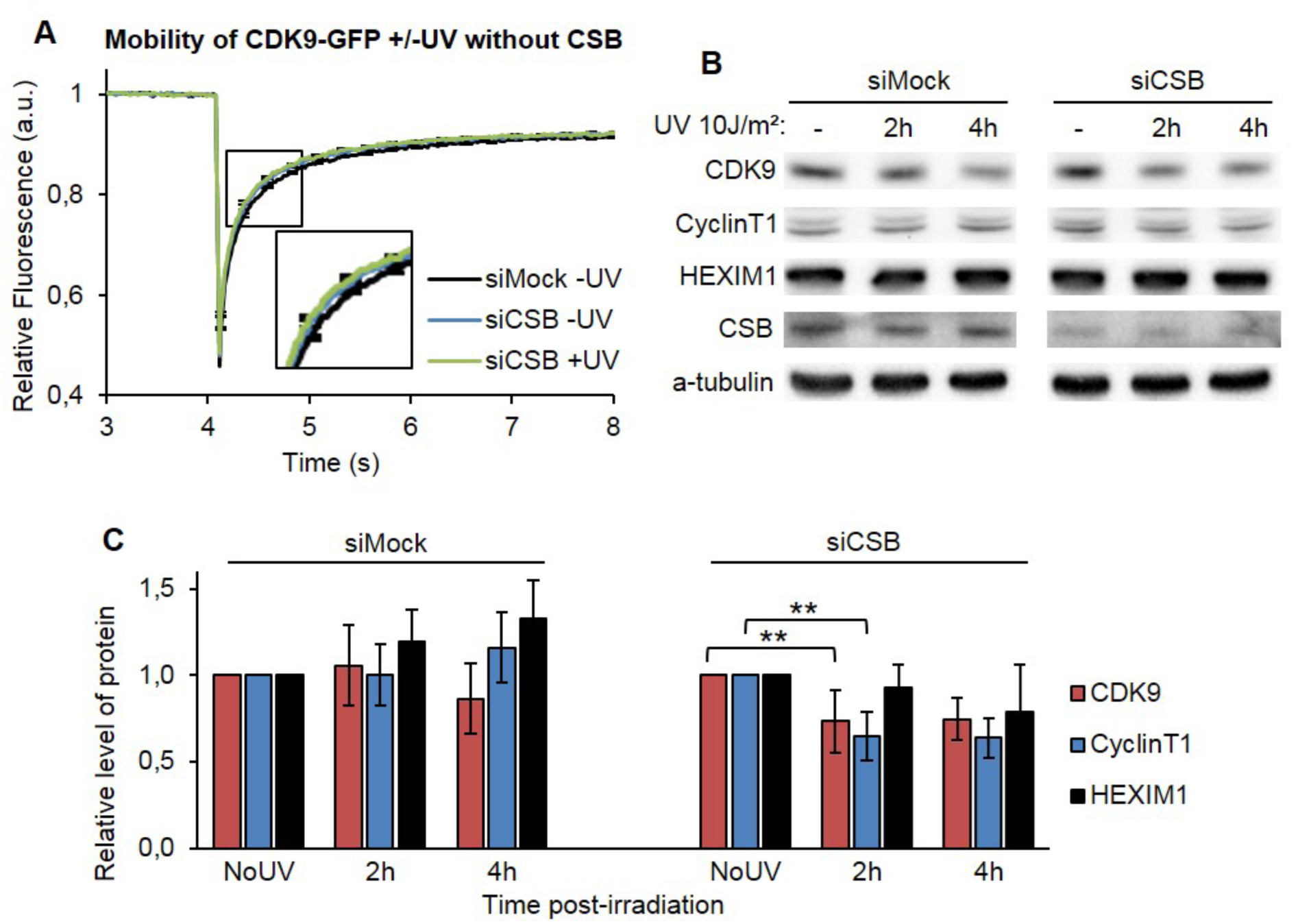
CDK9/CyclinT1 degradation without CSB. **A** Normalized FRAP analysis of CDK9-GFP expressing cells after siRNA mediated knockdown of the indicated factors. Cells were measured untreated (-UV) or 4h after UC-C exposure (+UV). Error bars represent the SEM obtained from at least 27 cells from 2 independent experiments. **A** Immunoblot analysis of CDK9, CyclinT1, HEXIM1 and CSB in MRC5 cells treated with siRNA mediated knockdown of the indicated factors followed by different time of irradiation. **B** Quantification of western blot. The values of each protein are normalized to no-UV condition.

These combined results (FRAP and Immuno-blot) demonstrate that specifically after UV irradiation the complex CDK9/CyclinT1 is released from Hexim1 complex, but if the DNA repair factor CSB is absent, this free fraction is rapidly degraded after UV irradiation.

### UV irradiation induces a CSB-dependent RNAP2 Serine 2 phosphorylation

During transcription-coupled repair, the first protein that encounters the lesion is the RNAP2. Not capable of bypassing the UV-lesion, the RNAP2 is stalled or backtracked and paused. This transcriptional arrest triggers the TCR reaction. When DNA repair is efficiently performed, RNAP2 will be released from this arrest and transcription will restart. Because this process is highly evocative of RNAP2 pausing downstream of promoters and CDK9 is involved in the phosphorylation and release of paused RNAP2 (20), we wanted to investigate whether during TCR, RNAP2 was specifically phosphorylated. We examined Serine2, Serine5 and Serine7 RNAP2 phosphorylation (21, 22) after UV irradiation in WT cells, 2 and 4 hours after UV-irradiation by western blot of nuclear extract, in this way no crosslink, which could interfere with the accessibility of the epitope (23), has been performed. While there was no difference in Serine5 and Serine7 RNAP2 phosphorylation after UV irradiation, a well-defined increase in Serine2 RNAP2 phosphorylation was observed 2 hours after UV-damage induction (Fig 4A). Interestingly, in CSB knocked down cells, this specific Ser2 RNAP2 phosphorylation was abolished (Fig 4B and 4C), showing that this phosphorylation is indeed specific for TCR reactions. Surprisingly, CDK9 depleted cells presented a normal Ser2 RNAP2 phosphorylation 2 hours after UV-irradiation (Fig 4D and 4E), demonstrating that CDK9 is not the kinase responsible for this phosphorylation. Because CDK12 can replace CDK9 for Ser2 phosphorylation (16, 24), we investigated whether the kinase activity of CDK12 could be responsible for this specific TCR Ser2 RNAP2 phosphorylation. CDK12 knocked down cells were irradiated and Ser2 RNAP2 phosphorylation was quantified 2 hours after UV. Surprisingly, a reduction of CDK12 does not affect this TCR-specific RNAP2 phosphorylation (Fig 4F and 4G). To confirm this result we quantified the TCR-dependent Ser2 RNAP2 phosphorylation in cells depleted for the different Cyclins associated with both CDK9 and CDK12: Cyclin K, CyclinT1, Cyclin T2, Cyclins T1/T2. Our results show that none of these Cyclins are associated with the TCR-specific Ser2 RNAP2 phosphorylation (Fig S5). Obviously, also in these experiments quantifications were normalized to the undamaged condition to be able to highlight exclusively the role of these kinases and cyclines after UV irradiation during the TCR reaction.

**Figure 4.**
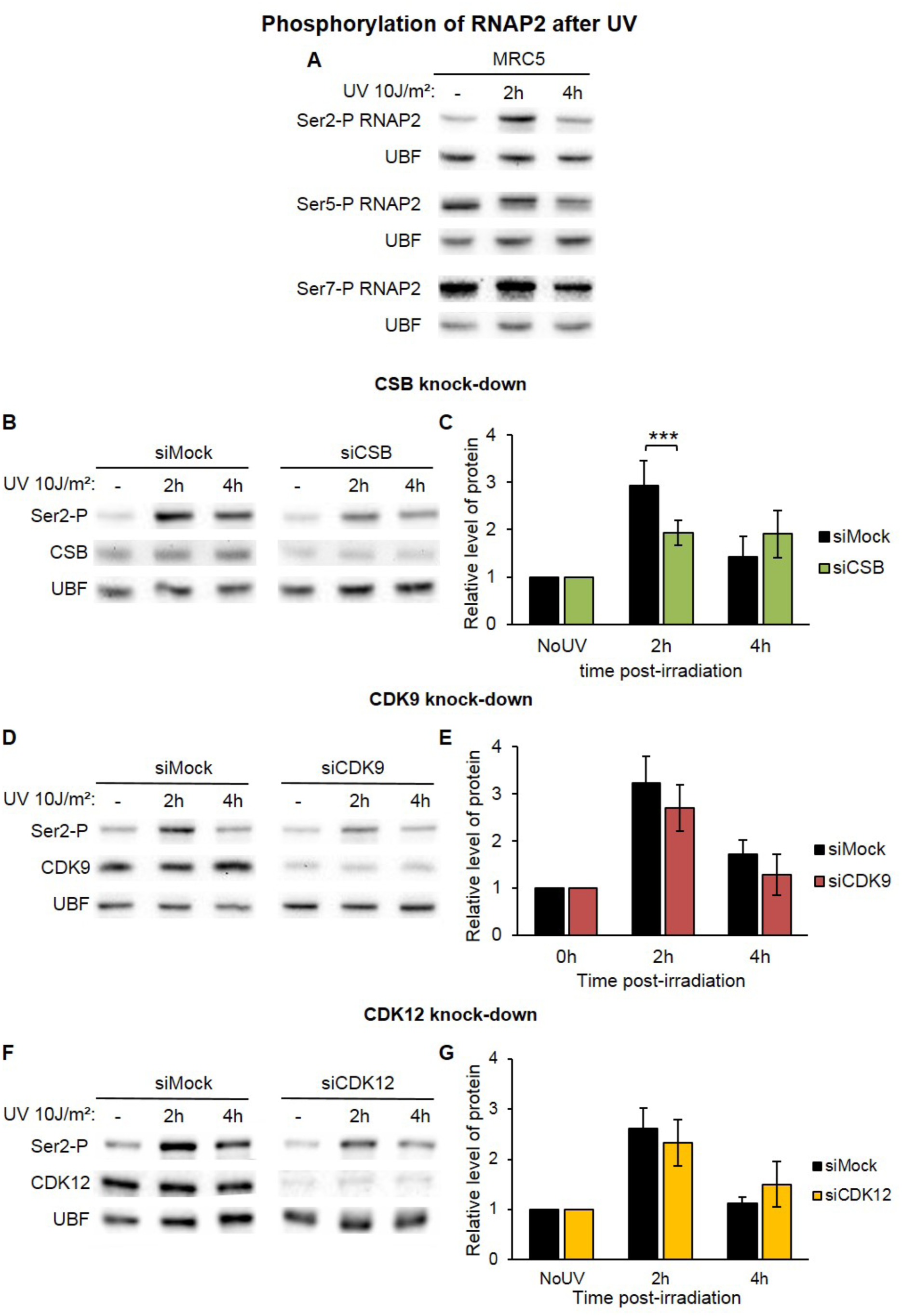
RNAP2 behavior in transcription restart after DNA damage. **A** Immunoblot showing the phosphorylation of Serine 2, 5 or 7 of RNAP2 in MRC5 whole cell extract after UV irradiation. **B-D-F** Western blot showing the phosphorylation of Serine 2 in MRC5 nuclear extract treated or not with **UV-C** after siRNA mediated knockdown of indicated factors showing the phosphorylation of Serine 2. UBF serves as a loading control. **C-E-G** Quantification of at least three different experiments of the RNAP2 phosphorylation/UBF ratio compared with NoUV condition for each siRNA. p-value: ^**^<0,05.

### CDK9 increases the mobility of RNAP2 after UV-irradiation

Because the kinase activity of CDK9 is not involved in the phosphorylation of RNAP2 during TCR, but having established that CDK9 plays a role in RTR (Fig 1), we planned to investigate whether the absence of CDK9 would affect the mobility of the RNAP2 during the repair reaction. In order to answer this question, we produced a plasmid expressing a GFP-tagged version of RNAP2 (Fig S6A) and stably transfected WT cells. We could show that the GFP-Pol2 is localized in the nucleus of cells and excluded from the nucleolus, as endogenous RNAP2 (Fig S6B). The GFP-Pol2 fusion protein is overexpressed in the overall population of cells (Fig S6C) and for this reason just low expressing cells were chosen to perform the FRAP analysis (Fig S6D). Strip-FRAP assays showed that GFP-Pol2 is largely immobilized, showing only a limited fraction of recovered protein after the photo-bleach. This immobile fraction represents the RNAP2 molecules engaged in the process of transcription and therefore retained on the chromatin. In order to verify this transcriptional engagement, we measured the mobility of RNAP2 after treatment with DRB (5,6-Dichloro-1-β-D-ribofuranosylbenzimidazole), an inhibitor of RNAP2 transcription. After treatment, the immobile fraction of GFP-Pol2 was reduced, indicating that the majority of GFP-Pol2 molecules are engaged towards the transcription process (Fig S6E).

To investigate the mobility of RNAP2 during RTR, we carried out Strip-FRAP experiments on GFP-Pol2 expressing cells in presence or absence of UV damage. As already observed in a previous study in our group (8), there is no measurable change of the mobility of GFP-Pol2, after UV damage (Fig 5A). This is explained by the fact that the majority of the RNAP2 immobile fraction is due to GFP-Pol2 molecules involved in transcription and that no additional measurable immobile fraction comes from lesion-stalled RNAP2 molecules.

However, in the absence of CDK9 and in the absence of DNA damage, the RNAP2 immobile fraction increases (Fig 5B). This result is probably due to the higher retention of RNAP2 molecules on transcriptional paused sites at proximal promoters, since CDK9 is essential for the RNAP2 release of paused sites and RNAP2 engagement in productive transcription elongation (25). The absence of CDK9, in combination with UV damage, results in an even more significant RNAP2 immobile fraction (Fig 5C), indicating that there are more RNAP2 molecules retained on the chromatin (molecules stalled on UV-lesions), additionally to the ones observed by default at the pausing sites.

**Figure 5.**
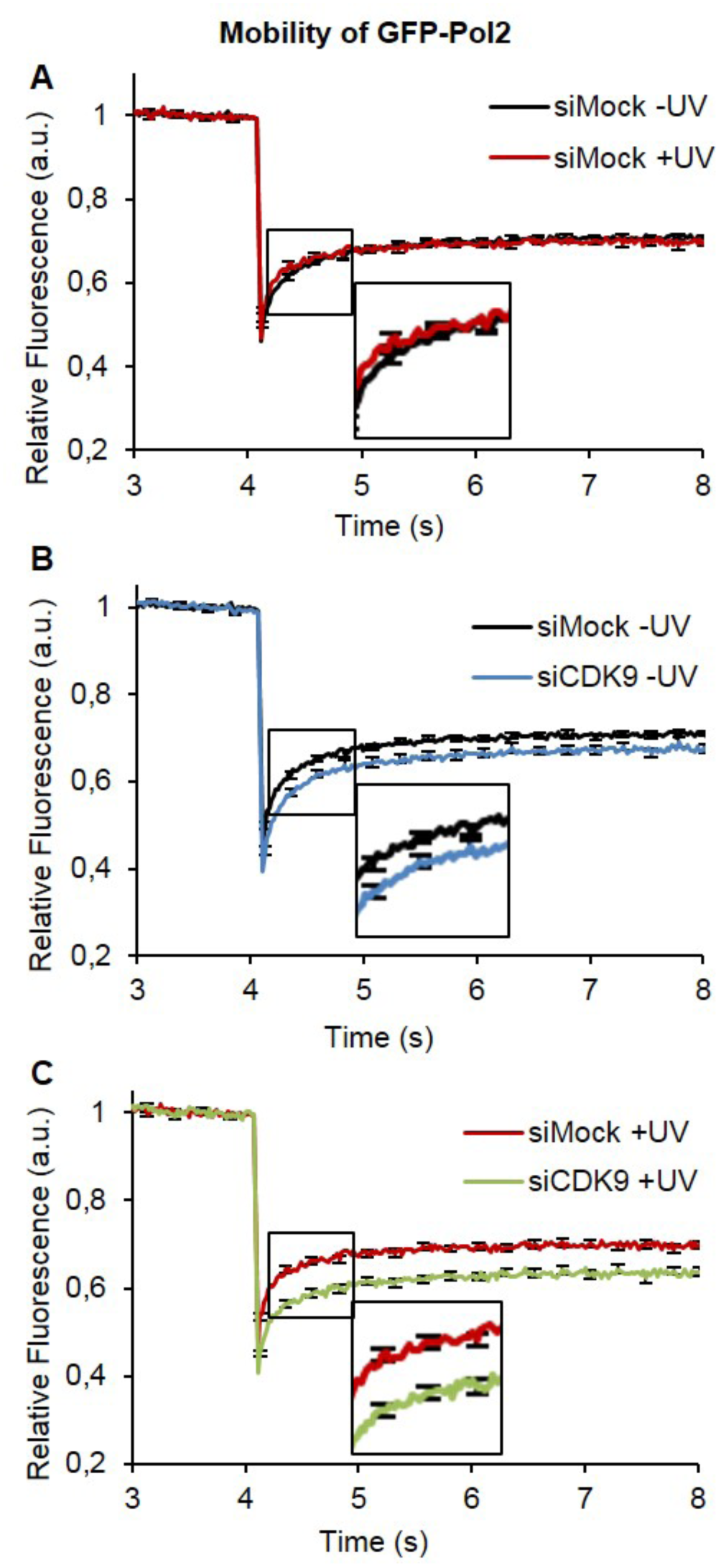
CDK9 function in transcription restart after UV damage repair. Normalized FRAP analysis of GFP-Pol2 expressing cells after siRNA mediated knockdown of indicated factors. Cells were measured untreated (-UV) or 4h after UC-C exposure (+UV). Error bars represent the SEM obtained from at least 27 cells from 2 independent experiments. p-value: siMock –UV and siCDK9 –UV < 0,01, siMock +UV and siCDK9 +UV < 0,001.

## Discussion

Bulky DNA-damage distorting the DNA helix, challenges constantly cell survival by interfering and blocking cellular functions, such as replication and transcription. During evolution, cells have developed processes that counteract the deleterious effect of these damages, restoring an undamaged DNA molecule and allowing the restart of cellular processes. The importance of rapidly restoring cellular functions is better demonstrated by the existence of DNA repair processes tightly coupled with RNAP2 transcription. When bulky DNA damages, such as ultraviolet lesions, are located on the transcribed strand of active genes, RNAP2 is stalled and transcription elongation is blocked. This pausing of transcription is necessary to allow the repair machineries to be recruited on the site of damage for the repair reaction to take place. Once the repair is completed, RNAP2 pausing may be released and transcription may restart.

During transcription and in absence of any DNA damage, RNAP2 naturally pauses at promoter-proximal sites, 30 to 60 nucleotides downstream of transcription start site (16). The release of this pausing into productive transcription elongation is under the control of P-TEFb, a heterodimeric cyclin-dependent kinase composed of CDK9 and CycT1/CycT2 (26). Because CDK9 interacts tightly with ELL and it has been demonstrated that ELL is involved in the RTR and is recruited to the damage via CdK7 (8), we wondered whether CDK9 could play a role in RTR and more importantly, if RTR would share a common molecular mechanism with the release of RNAP2 promoter-proximal pausing. In this article, our effort was focused on decrypting the role of CDK9 during RTR, since as well as for ELL (8), reduction of CDK9 concentration in CDK9-knocked down cells severely delays the RTR process without affecting the repair reaction *per se* (Fig1). In order to exclusively investigate the effect of CDK9 reduction during TCR, we always normalized our results to the undamaged condition in which we measured that a 30% CDK9 reduction (obtained with several combinations of siRNAs) results in a 70% reduction of basal transcription activity. To investigate CDK9 action on damaged chromatin, we produced a GFP-labeled version of CDK9 and we quantified the mobility of CDK9-GFP before and after UV by Fluorescence Recovery After Photo-bleaching (FRAP) assays. During Nucleotide Excision Repair, repair proteins are partially and temporally immobilized on the damaged site and this dynamic behavior influences the FRAP curves before and after UV irradiation. Namely, repair proteins mobility is slower after UV damage XPA (27). ELL was found to behave as a canonical repair protein, and the immobile fraction observed after DNA repair as well as the immobilization on a locally damaged area was comparable to the one measured for TCR-specific proteins, such as CSA (8). Surprisingly, CDK9 presented a different dynamic behavior after UV irradiation, instead of being more immobile, FRAP curves pointed towards a remobilization of the protein. More remarkably, this remobilization was CSB-dependent. To be able to fine-tune RNAP2 pause release, CDK9 kinase activity is kept under tight control by its interaction with a large inhibiting complex (7SK snRNP)(26). Indeed, activation of the CDK9/CycT implies a release from 7SK snRNP and in particular the dissociation from the P-TEFb inhibitor protein HEXIM1 (18). After UV irradiation, the increased mobility measured by FRAP analysis for CDK9 is concomitant with the dissociation of CDK9/CycT from HEXIM1 (Fig 2 and Fig 6). Surprisingly, after UV-irradiation in CSB mutant cells and CSB knocked-down cells, cellular CDK9-CycT amount is reduced. This novel observation points to a possible direct or indirect role of CSB in stabilizing the CDK9/CycT complex on the damaged site. In the absence of CSB, this stabilization is likely not achievable and the CDK9/CycT complex is degraded (Fig 3 and Fig 6C, after UV without CSB). During TCR, it has been proposed that RNAP2 would backtrack and remain in the proximity of the damaged site to restart transcription or that RNAP2 would detach from chromatin and restart transcription from the promoter (28). It is not excluded that these processes could in fact coexist depending on the site of damage (i.e. RNAP2 stalled at a lesion proximal to initiation or termination would be more prone to disengage from the DNA) or the time it takes for the repair (i.e. longer repair timing could lead to RNAP2 disengagement). In both cases (at the site of damage or at the promoter), RNAP2 needs to restart the transcription and this step requires phosphorylation of the CTD at one of the Serines (2, 5 or 7). We could show here that, after UV-irradiation, hence during TCR-RTR, RNAP2 is strongly phosphorylated at Ser2. This Ser2 phosphorylation is specifically observed 2 hours after UV irradiation and is less abundant at 4 hours after irradiation. More interestingly, this Ser2 phosphorylation is specifically observed in TCR-proficient cells but absent, or greatly reduced, in TCR-deficient cells (Fig 4), demonstrating that RNAP2 Ser2 phosphorylation is the one specifically needed for the restart of transcription after completion of repair. Surprisingly, we could demonstrate that CDK9 (nor CDK12) is not the kinase that is in charge for this phosphorylation, which still takes place in CDK9 depleted cells (Fig 4 and Fig 6). This result was unexpected as it shows that the complex CDK9/CycT plays a different role in RTR than it has in the release of pausing at proximal promoters.

**Figure 6.**
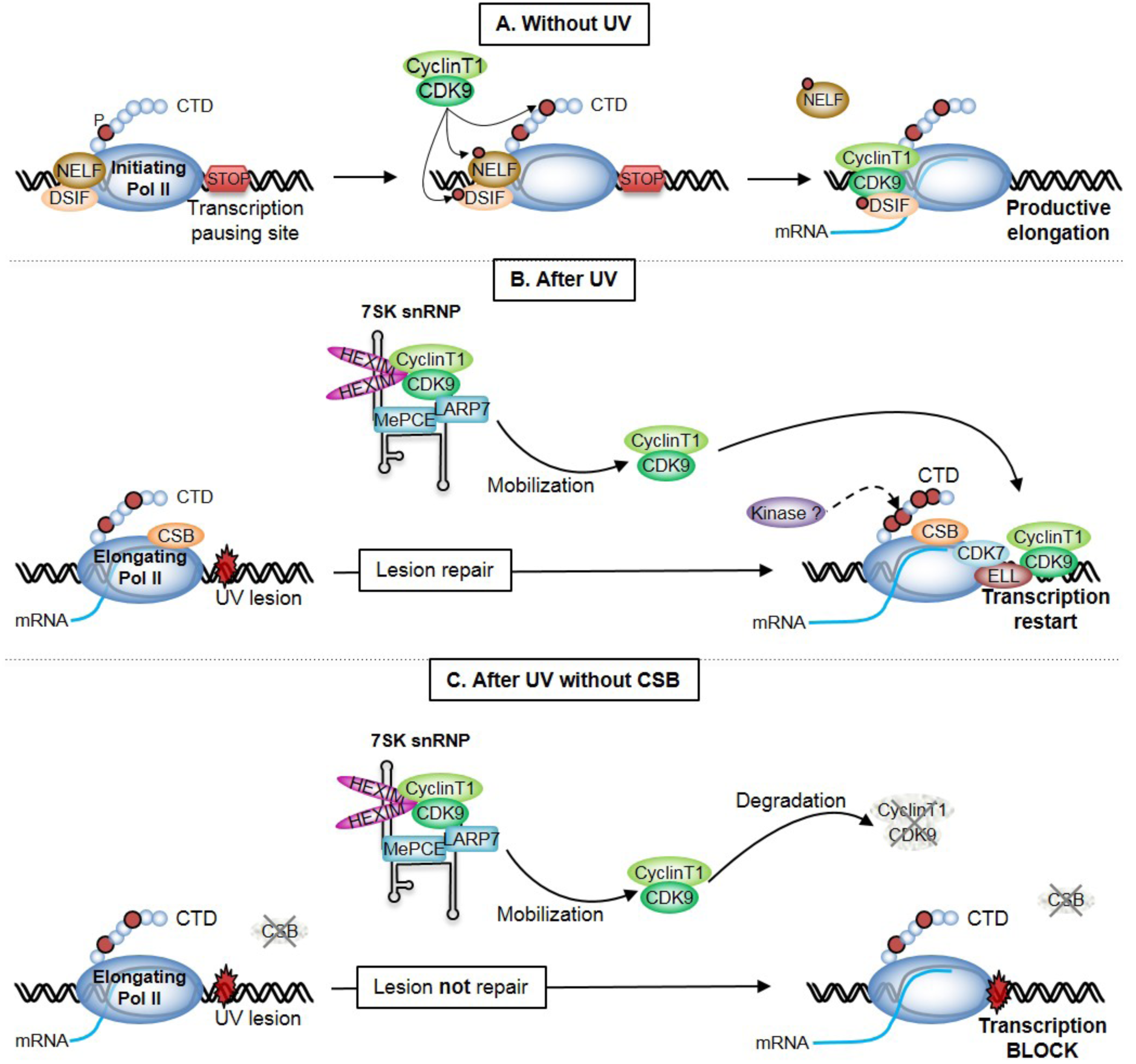
Mechanistic model of CDK9’s role in transcription restart after DNA repair. **A.** Immediately following initiation, RNAP2 soon enters transcriptional arrest mediated by DSIF and NELF. Their negative effect can be relieved by P-TEFb composed of CDK9 and CyclinT. The kinase CDK9 phosphorylates (red circle) DSIF and NELF as well as the serine 2 of RNAP2 CTD allowing productive elongation. **B.** After UV irradiation, CDK9 and CyclinT is released from 7SK snRNP complex and can be recruited to the site of damage probably via its interaction with ELL. After completion of the repair reaction (lesion removal and DNA gap filing), CDK9 plays an essential role to enable RNAP2 restart, probably as a docking protein to recruit additional factors needed for transcription resumption. **C.** Without CSB, lesion is not repaired. CDK9/CyclinT1 is release from 7SK snRNP complex and without CSB is probably not recruited to the site of damage leading to a rapid degradation of the CDK9/CyclinT1. Transcription remains blocked.

Because CDK9 reduction has a clear effect on the timing of the restart of transcription after DNA repair achievement (Fig 1) but has no kinase activity on RNAP2 (Fig 4), we explored the effect of the absence of CDK9 on the mobility of RNAP2 molecules. In CDK9 knocked down cells and without any damage, more RNAP2 molecules are binding to the chromatin. This result is in accordance with the function of CDK9 during release of pausing, i.e. in its absence more RNAP2 molecules are paused on proximal promoters. However, after UV irradiation, an even further RNAP2 immobile fraction is observed in CDK9 knocked down cells, supporting the hypothesis that without CDK9, RNAP2 is more bound to the damaged chromatin and that the release of RNAP2 is impeded or at least retarded (Fig 5). Interestingly, in our previous study, we observed that this increased RNAP2 immobile fraction is also observed when cells are ELL-depleted (8). This result could suggest that ELL and CDK9 would play a structural role during RTR, facilitating other proteins to be either recruited or stimulated in their function of restarting transcription after completion of DNA repair. In conclusion, the role and mechanism of action of CDK9 is specific during RTR (Fig 6B, after UV) and is different from its function in the release of paused RNAP2 (Fig 6A, without UV), indicating that although RTR and pausing might share common actors, the mechanism of action of RTR is unexpected and remains to be fully explored, starting from finding the kinase that is responsible for the specific RNAP2 Ser2 phosphorylation.

## Materials and Methods

### Cell culture

Cell strains used were: (*i*) wild type SV40-immortalized human fibroblasts (MRC5-SV); (*ii*) MRC5-SV stably expressing GFP-Pol II (G418 selected-0.2 mg/ml); (*iii*) MRC5-SV stably expressing CDK9-GFP (G418 selected-0.2 mg/ml); (*iv*) CSB deficient SV40-immortalized human fibroblasts (CS1AN, TCR-deficient); (v) MRC5-SV stably expressing CSA-GFP (G418 selected-0.2 mg/ml) (*vi*) XPC deficient SV40-immortalized human fibroblasts (XP4PA, GGR-deficient). Human fibroblasts were cultured in a 1:1 mixture of Ham’s F10 and DMEM (Lonza) supplemented with antibiotics (penicillin and streptomycin) and 10% fetal calf serum, at 37°C and 5% CO_2_.

### Specific treatment

DNA damage was inflicted by UV-C light (254 nm, 6W lamp). For UV survival experiments, cells were exposed to different UV-C doses, 1 day after plating. Survival was determined by clone counting, 10 days after UV irradiation, as described previously (29). For FRAP, RRS, and UDS experiments, cells were globally irradiated with 16 J/m^2^ of UV-C and 10J/m^2^ for immunoblot experiment whereas for TCR-UDS, cells were locally irradiated with 100 J/m^2^ m of UV-C through a 5-μm-pore polycarbonate membrane filter (Millipore).

GFP-Pol2 expressing cells were treated with 100 μg/ml of 5,6-dichloro-1-beta-D-ribofuranosylbenzimidazole (DRB, sigma) for 2h at 37°C before FRAP analysis.

### Construction and expression of GFP-Pol 2 and CDK9-GFP fusion protein

Full length RNAP2 c-DNA was cloned in-frame into pEGFP-C1 vector and full length CDK9 c-DNA was cloned in-frame into pEGFP-N1 vector (Clontech). Constructs were sequenced prior to transfection. Transfection in MRC5-SV40 transformed human fibroblasts was performed using Fugene transfection reagent (Roche). Stably expressing cells were isolated after selection with G418 (Gibco) and single cell sorting using FACS (FACScalibur, Beckton Dickinson).

### Fluorescence Recovery after Photobleaching (FRAP)

FRAP experiments were performed as described before (17) on a Zeiss LSM 710 NLO confocal laser scanning microscope (Zeiss), using a 40x/1.3 oil objective, under a controlled environment (37°C, 5% CO_2_). Briefly, a narrow region of interest (ROI) centred across the nucleus of a living cell was monitored every 20 (1% laser intensity of the 488 nm line of a 25 mW Argon laser) until the fluorescence signal reached a steady state level (after circa 2 s). The same strip was then photobleached for 20 ms at 100% laser intensity. Recovery of fluorescence in the strip was then monitored (1% laser intensity) every 20 for about 20 seconds. Analysis of raw data was performed with the ZEN software (Zeiss). All FRAP data were normalized to the average pre-bleached fluorescence after background removal. Every plotted FRAP curve is an average of at least twenty measured cells.

### Laser micro-irradiation

In order to locally induce DNA damage in living cells, we used a tuneable near-infrared pulsed laser (Cameleon Vision II, Coherent Inc.) directly coupled to an inverted confocal microscope equipped with a 40x/1.3 oil objective and a thermostatic chamber maintained at 37°C with 5% CO_2_ (LSM 710 NLO, Zeiss). Typically, a small circular area (3 μm in diameter) within the nucleus of a living cell was targeted 3 times to 15% of the laser (800 nm). Subsequent time-lapse imaging of targeted cells was performed every 15s for 420s. Image analysis was performed using ImageJ (Rasband, W.S., National Institutes of Health, USA) and a custom-built macro as follows: (i) the time series image stack was adjusted to compensate for cell movement (StackReg plugin), (ii) a ROI spanning the total nucleus was defined to compensate for unwanted photobleaching during the acquisition of images, (iii) a ‘local damage’ ROI was specified to quantify the fluorescence increase due to (GFP tagged) protein recruitment at the laser induced DNA damage area. At least ten cells were measured for all cell lines.

### RNA interference

Short interfering RNAs (siRNAs) used in this study are pool of siRNA and are: siMock, Dharmacon D-001810-01 (10 nM); siCDK9, Dharmacon L-003243-00 (10 nM); siCDK12, Sanatcruz sc-44343 (5 nM); siCSB, Dharmacon L-004888-00 (10 nM); siCyclinK, Dharmacon L-029590-00 (10 nM); siCyclinT1, Dharmacon L-003220-00 (10 nM); siCyclinT2, Dharmacon L-003221-00 (10 nM); siXPF, Dharmacon M-019946-00 (10 nM). In parenthesis, the final concentration used for each siRNA. Cells were transfected with siRNA using GenJET siRNA transfection reagent (Tebu-Bio) according to the manufacturer’s protocol. Briefly, 100.000 cells were seeded per wells of a 6-wells plates and allowed to attach overnight. Transfection complexes were formed by 15 min incubation at room temperature using buffer provided and added 24h after seeding. A second siRNA transfection was performed 24h after the 1^st^ and cells were grown confluent. Experiments were carried out 48h after the 1^st^ siRNA transfection. Proteins knock down was confirmed by western blot.

### Protein extraction

For protein extraction, cells were cultured either in a 10-cm dishes or in 6-wells plates if cells need to be transfected with siRNA. After irradiation as described above, cells were harvested by Trypsination. The extraction of proteins has been performed by using either the kit CelLytic^™^ NuCLEAR^™^ Extraction (Sigma-Aldrich) for nuclear extract or the kit Mammalian Cell Lysis Reagent^™^ (Sigma-Aldrich) for total protein extraction. The concentration of proteins has been determined by using the Bradford method. Then, samples were diluted with 2X Laemmli buffer, heated at 95°C and loaded on a SDS-PAGE gel.

### Co-immunoprecipitation

For co-immunoprecipitation, 10μl of protein G magnetic bead (Bio-adembead, Ademtech) were used par IP. 1μg of anti-HEXIM1 antibodies (rabbit, A303-113A, Béthyl) were bound to the beads in PBS with BSA (5mg/ml) during 2h at 4°C with rotation. 200μg of whole cell extract were then incubated with beads-antibodies complex for 2h at 4°C with rotation. After 2 washes at 100mM salt, 2 washes at 150mM and 1 wash at 100mM, beads were boiled in 2x Laemmli buffer and loaded on a SDS-PAGE gel.

### SDS-PAGE

Proteins were separated on SDS-PAGE composed of bisacrylamide (37:5:1), blotted onto a polyvinylidene difluoride membrane (PVDF, 0.45μm Millipore) and analysed using the following primary antibodies: anti-Serine2 phosphorylation RNAP2 (rabbit, ab5095 Abcam); anti-Serine5 phosphorylation RNAP2 (rabbit, #13523, cell signalling); anti-Serine7 phosphorylation RNAP2 (rabbit, #13780, cell signalling); anti-RNAP2 (rabbit, sc-899 Santa Cruz Biotechnology); anti-CDK9 (rabbit, sc-8338X Santa Cruz Biotechnology); anti-CSB (goat, sc10459 Santa Cruz Biotechnology); anti-CyclinK (rabbit, ab57311, abcam); anti-CyclinT1 (rabbit, sc-10750 Santa Cruz Biotechnology); anti-CyclinT2 (mouse, ab50979, abcam); anti-HEXIM1 (rabbit, A303-113A, Béthyl); anti-α-tubulin (mouse, T6074, sigma-aldrich); anti-UBF (mouse, sc-13125 Santa Cruz Biotechnology) and anti-TBP (mouse, 3TF1-3G3, ThermoFisher). The loading was controlled with either the anti-UBF, anti-TPB or anti-α-tubulin antibody. Western blot process was performed as described previously (Rockx et al. 2000) and protein bands were visualised via chemiluminescence (ECL <u>E</u>nhanced <u>C</u>hemo <u>L</u>uminescence; Pierce ECL Western Blotting Substrate) using Horseradish Peroxidase (HRP)-conjugated secondary antibodies and imaged via the chemidoc system (BioRad). The quantification of the band was performed with the software Image Lab (BioRad) using the method of volumes (rectangle). The background was removed with the Local subtraction method.

### Recovery of RNA synthesis (RRS) assays

MRC5-SV40 cells were grown on 24 mm coverslips. siRNA (siCDK9/siCSB) transfections were performed 24h before RRS assays. RNA detection was performed using a Click-iT RNA Alexa Fluor Imaging kit (Invitrogen), according to the manufacturer’s instructions. Briefly, cells were UV-C irradiated (16 J/m^2^) and incubated for 0, 3, 16 and 24 h at 37°C. Then, cells were incubated for 2 hours with 5-ethynyl uridine, fixed and permeabilized. Cells were incubated for 30 min with the Click-iT reaction cocktail containing Alexa Fluor Azide 488. After washing, the coverslips were mounted with Vectashield (Vector). Images of the cells were obtained with the same setup (see FRAP methods section) and constant acquisition parameters, then the average fluorescence intensity per nucleus was estimated after background subtraction (using ImageJ) and normalized to not treated cells. For each sample, at least 80 nuclei were analysed from three independent experiments.

### Unscheduled DNA synthesis (UDS) assays

MRC5-SV40 cells were grown on 24 mm coverslips. siRNA (siCDK9/siXPF) transfections were performed 24h before UDS assays. *De novo* DNA synthesis detection was performed using a Click-iT DNA Alexa Fluor Imaging kit (Invitrogen), according to the manufacturer’s instructions. Briefly, after global irradiation cells were incubated for 3 hours with 5-ethynyl-2’-deoxyuridine (EdU), then cells are washed with PBS, fixed and permeabilized. Fixed cells were incubated for 30 min with the Click-iT reaction cocktail containing Alexa Fluor Azide 594. After washing, the coverslips were mounted with Vectashield containing DAPI (Vector). Images of the cells were acquired being careful not to take cells in replication. Images were analysed in the same way as for the RRS assay (see previous paragraph). For each sample, at least 20 nuclei were analysed from three independent experiments.

### TCR-UDS assays: UDS measurement during TCR

XPC deficient SV40-immortalized human fibroblasts (XP4PA-GGR deficient cell line), were grown on 24 mm coverslips. siRNA (siCDK9/siXPF) transfections were performed 24h before UDS assays. After local irradiation (100 J/m^2^ UV-C) through a 5 μm pore polycarbonate membrane filter, cells were incubated for 8 hours with ethynyldeoxyuridine, washed, fixed and permeabilized. Fixed cells were treated with a PBS-blocking solution (PBS+: PBS containing 0.15% glycine and 0.5% bovine serum albumin) for 30 min, subsequently incubated with primary antibodies mouse monoclonal anti-yH2AX (Ser139) (Upstate, clone JBW301) 1/500 diluted in PBS+ for 1h, followed by extensive washes with Tween20 in PBS. Cells were then incubated for 1h with secondary antibodies conjugated with Alexa Fluor 488 fluorescent dyes (Molecular Probes, 1:400 dilution in PBS+). Then, cells were incubated for 30 min with the Click-iT reaction cocktail containing Alexa Fluor Azide 594. After washing, the coverslips were mounted with Vectashield containing DAPI (Vector). Images of the cells were obtained with the same microscopy system and constant acquisition parameters. Images were analysed using ImageJ as follows: (i) a ROI outlining the locally damaged area was defined by using the yH2AX staining, (ii) a second ROI of comparable size was defined in the nucleus (avoiding nucleoli and other non-specific signals) to estimate background signal, (iii) the ‘local damage’ ROI was then used to measure the average fluorescence correlated to the EdU incorporation, which is an estimate of DNA replication after repair once the nuclear background signal obtained during step (ii) is subtracted. For each sample, between 50 and 60 nuclei were analysed from three independent experiments.

### Immunofluorescence

Cells were plated in 3.5 cm diameter wells on coverslips 24mm, in order to reach 70% confluency on the day of the staining. Cells were washed twice in PBS, fixed in 2% paraformaldehyde, permeabilized two times for 10min with PBS containing 0.1% Triton X-100 (PBS-T) and then washed with PBS containing 0.15% glycine and 0.5% bovine serum albumin (PBS+). We diluted antibodies in PBS+ and incubated cells with antibodies for 2 h at room temperature in a moist chamber. Antibodies used were: anti-CDK9 (rabbit, sc-484 Santa Cruz Biotechnology, 1/500 dilution), anti-RNA Pol II (rabbit, sc-899 Santa Cruz Biotechnology, 1/500 dilution). After washing (3 times short wash, 2 times for 10 min each with PBS-T) cells were incubated with the secondary antibody coupled to a fluorochrome: goat anti-rabbit conjugated with Alexa 488 (A11088 Invitrogen), 1/500 dilution in PBS+. After the same washing procedure, coverslips were mounted with Vectashield containing DAPI (Vector). Slides have been observed on a fluorescent microscope LSM710 NLO (Zeiss), using an objective 40x/1.3, and the analysis has been performed using ImageJ (NIH).

## Acknowledgments

We are grateful to Nicolas Heddebaut and Amandine Mourcet for the technical help provided.

This work was supported by l’Agence Nationale de la Recherche (ANR DyReCT: ANR-14-CE10-0009) and the ARC (Association pour la Recherche sur le Cancer) foundation (projet Fondation ARC PJA 20131200188).

L-M.D. and G.S. were supported by l’Agence Nationale de la Recherche (ANR DyReCT: ANR-14-CE10-0009). A.L. was supported by Association pour la Recherche sur le Cancer (Post-Doc grant). The funders had no role in study design, data collection and analysis, decision to publish, or preparation of the manuscript.

## ABBREVIATIONS

TCR: Transcription Coupled Repair
RTR: Resumption Of Transcription After Repair
NER: Nucleotide Excision Repair
RNAP2: RNA Polymerase II

---

**Figure S1.**
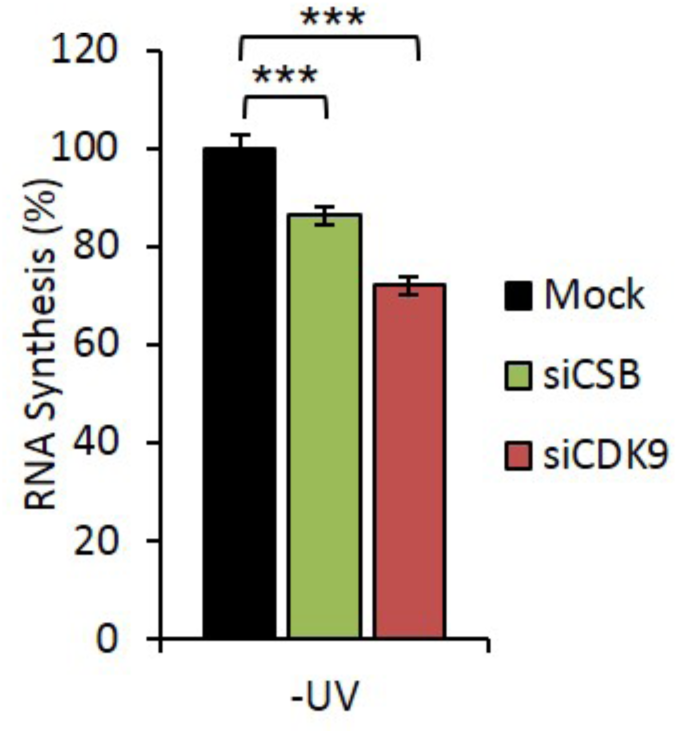
Level of RNA synthesis without UV. RNA synthesis in MRC5 cells after siRNA mediated knockdown of the indicated factors. At least 60 nuclei were analysed. Mock siRNA (black), CDK9 siRNA (red), CSB siRNA (green); error bars represent the SEM. p-value: ^***^<0,001.

**Figure S2.**
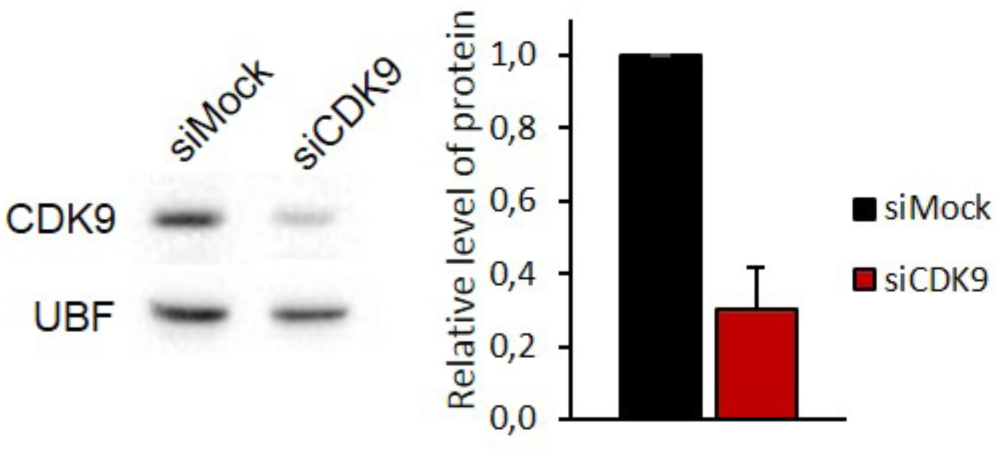
CDK9 siRNA efficiency. A Western blot showing the expression of CDK9 in MRC5 nuclear extract after siRNA mediated knock-down of CDK9. UBF serves as a loading control. B Quantification of at least three different experiments of CDK9 level normalized to UBF and siCDK9 compared to siMock.

**Figure S3.**
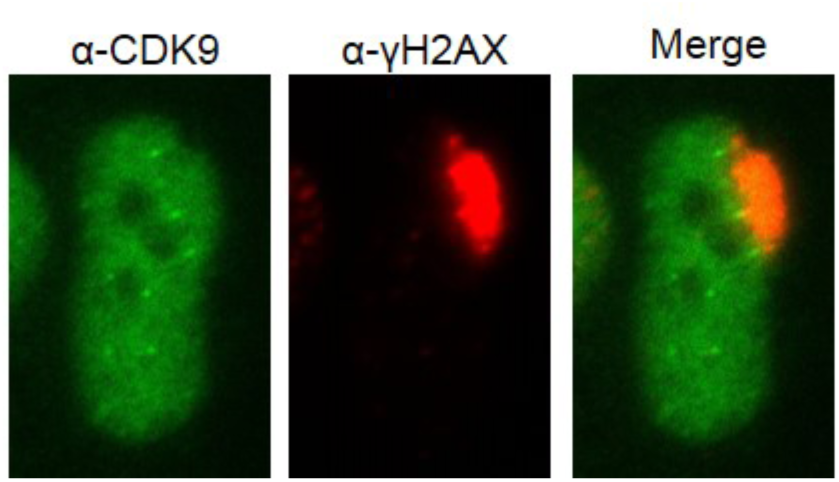
No recruitment of CDK9 on Local Damage. Immunofluorescence staining of MRC5sv cells with an anti-CDK9 and an anti-γH2AX antibody 3h after local damage induction with UV-C.

**Figure S4.**
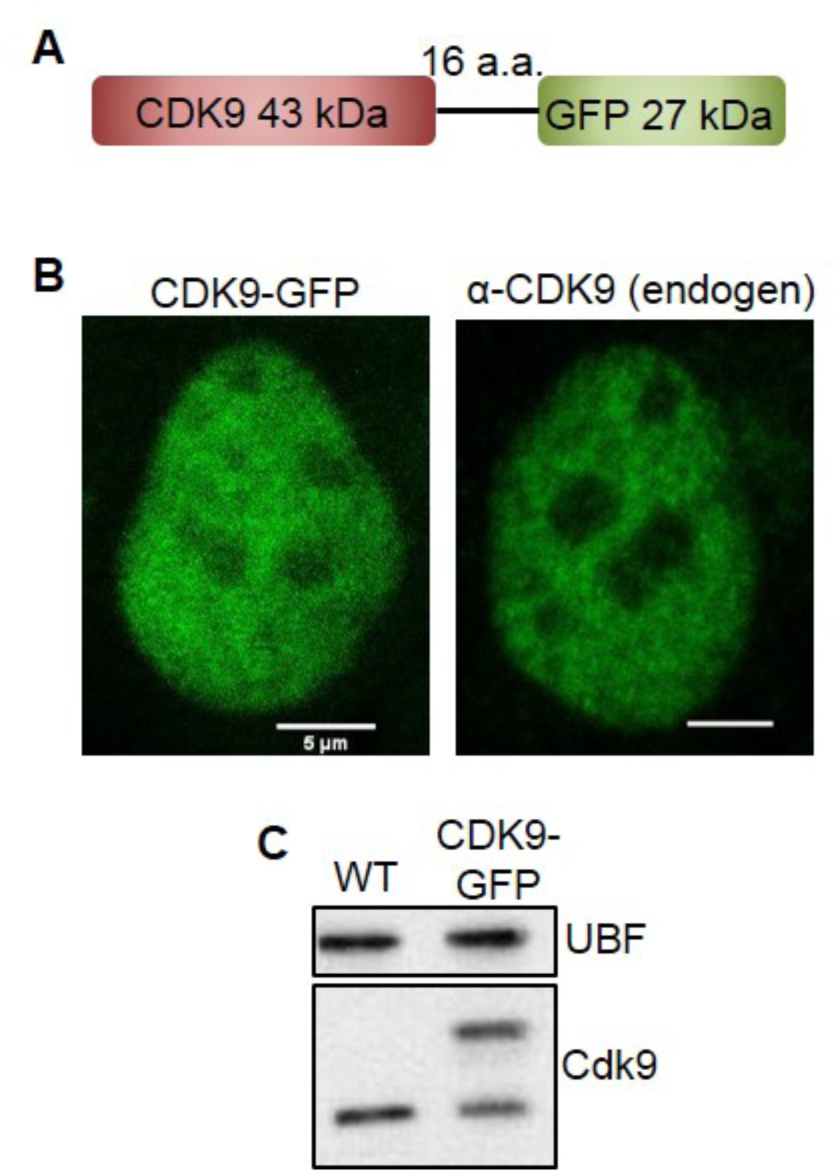
Characterization of stably expressing CDK9-GFP cell lines. **A** Scheme of the CDK9-GFP fusion protein **B** Confocal images of CDK9-GFP expressing cell lines and immunofluorescence staining of MRC5sv cells with a primary antibody against CDK9 **C** Immunoblot probed with an anti-CDK9 antibody of WT MRC5 cells and MRC5 cells stably expressing CDK9-GFP. UBF serves as a loading control.

**Figure S5.**
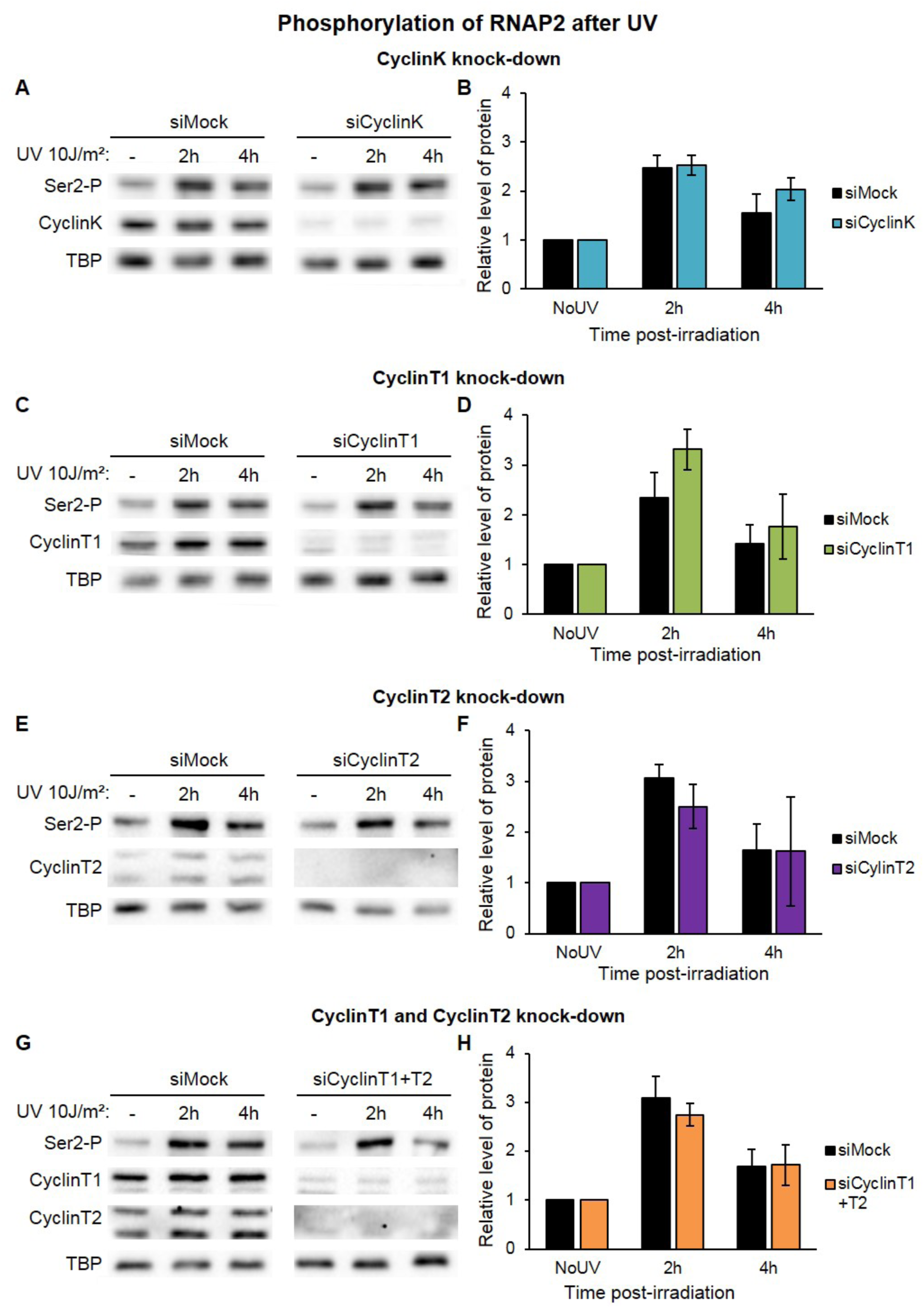
RNAP2 serine 2 phosphorylation without Cyclin proteins of CDK9. **A-C-E-G** Western blot showing the phosphorylation of Serine 2 in MRC5 nuclear extract treated or not with UV-C after siRNA mediated knockdown of indicated factors showing the phosphorylation of Serine 2. TBP serves as a loading control. **B-D-F-H** Quantification of at least two different experiments of the RNAP2 phosphorylation/TBP ratio compared with NoUV condition for each siRNA. p-value: ^**^<0,05.

**Figure S6.**
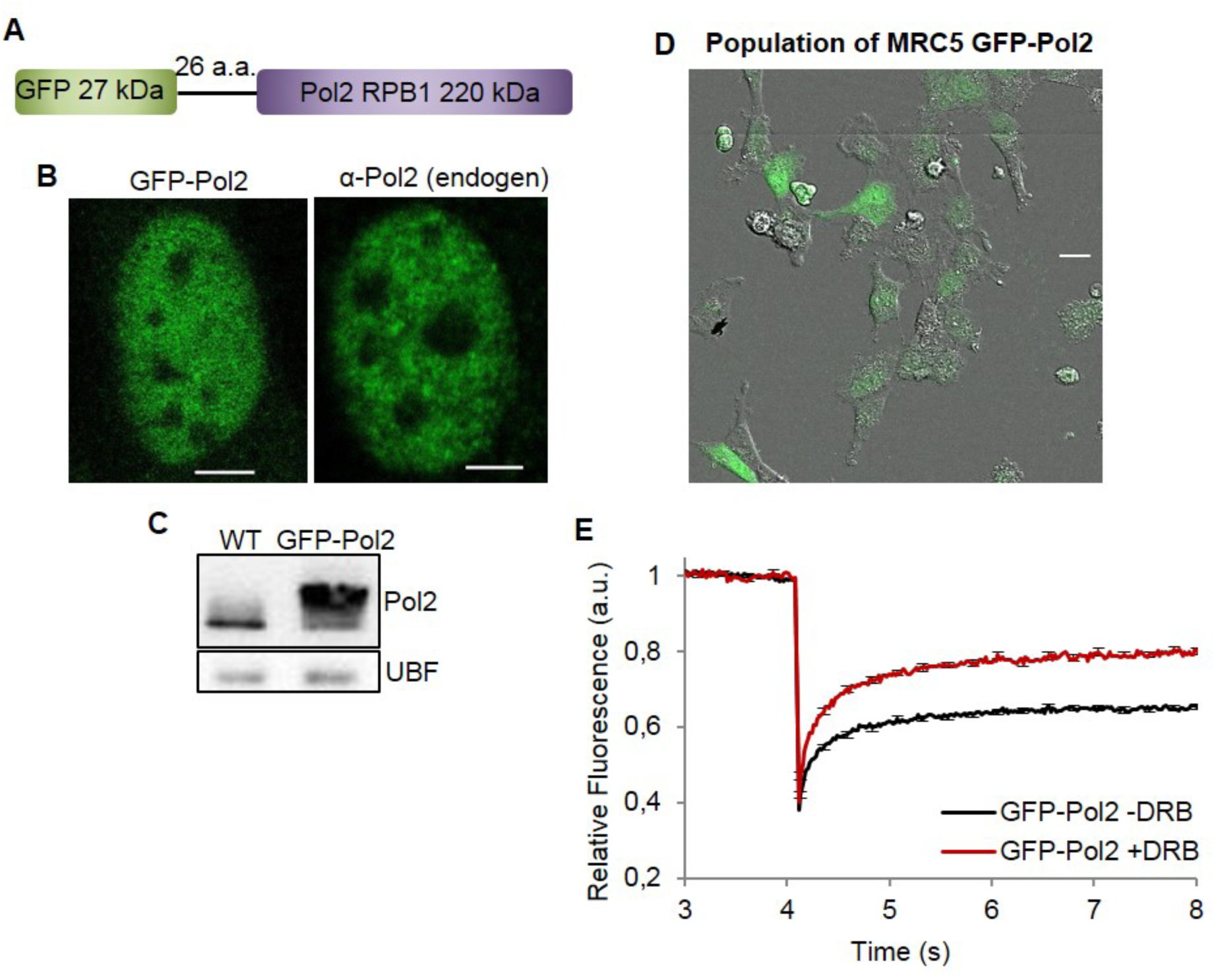
Characterization of the stably expressing GFP-pol2 cell lines. **A** Scheme of the GFP-Pol2 fusion proteins **B** Confocal images of GFP-Pol2 expressing cell lines and immunofluorescence staining of MRC5 cells with a primary antibody against total Pol2. Scale bar = 5μm **C** Immunoblot probed with an anti Pol2 antibody of WT MRC5 cells and MRC5 cells stably expressing GFP-Pol2. UBF serves as a loading control. **D** Confocal images of GFP-pol2 expressing cell lines. Scale bar = 20μm **E** FRAP analysis of GFP-Pol2 expressing cells untreated (red line) or treated with DRB (blue line). Error bars represent the SEM obtained from at least 15 cells.

